# Quantification of in-plane flexoelectricity in lipid bilayers

**DOI:** 10.1101/2021.01.29.428306

**Authors:** Nidhin Thomas, Ashutosh Agrawal

## Abstract

Lipid bilayers behave as 2D dielectric materials that undergo polarization and deformation in the presence of an electric field. This effect has been previously modeled by continuum theories which assume a polarization field oriented normal to the membrane surface. However, the molecular architecture of the lipids reveals that the heqadgroup dipoles are primarily oriented tangential to the membrane surface. Here, we perform atomistic and coarse-grained molecular dynamics simulations to quantify the in-plane polarization undergone by a flat bilayer and a spherical vesicle in the presence of an applied electric field. We use these predictions to compute an effective in-plane flexoelectric coefficient for four different lipid types. Our findings provide the first molecular proof of the in-plane polarization undergone by lipid bilayers and furnish the material parameter required to quantify membrane-electric field interactions.

## Introduction

Lipid membranes are 2D liquid crystal films that define the structural identity of cells and cellular organelles. Similar to 3D liquid crystals, lipids bilayers behave as dielectric materials that undergo polarization and deformation when subjected to electric fields. This electromechanical coupling forms the basis for various phenomena. For example, endogenous electric fields have been shown to play a critical role in would healing and tissue development [1–3]. Electric field-induced pore formation in bilayers is widely used to deliver genes and drugs into cells [4–8]. Electroporation is also employed to treat some cancers [9–11]. Electromechanical coupling plays a vital role in regulating the workings of auditory hair cells in mammals [12–14]. Electric field-induced vesicle deformation has been used to investigate membrane properties and stability [15–18].

Several theories have been proposed over the last two decades to model and comprehend the eletromechanical coupling in lipid bilayers, also referred to as flexoelectricity [19–24]. These frameworks have invariably invoked lipid dipoles and the resulting polarization field oriented normal to the membrane surface. However, lipid headgroup structure reveals that lipid dipoles predominantly lie in the tangential plane of the membrane surface. Figure 1 shows the dipole vector in a DPPC lipid. The dipole vector composed of the choline group and the phosphate group is oriented at 71° from the surface normal vector. As a result, the polarization field arising from the reorientation of the dipoles is expected to be primarily in the tangent plane. To incorporate this feature, Steigmann and Agrawal employed the 3D liquid crystal theory to derive a new electromechanical theory with in-plane polarization field [25]. This new model was recently used to predict shape transformations of confined vesicles [26]. While the application of the in-plane flexoelectric model has just begun, a quantification of the in-plane polarization and the required material parameter (in-plane flexoelectric coefficient) for the continuum model is still lacking. Here, in this study, we use atomistic studies and coarse-grained molecular dynamics to address this issue. Our simulations show that bilayers undergo significant in-plane polarization in the presence of a tangential electric field, yielding the material parameter for four lipid types.

**Fig. 1.**
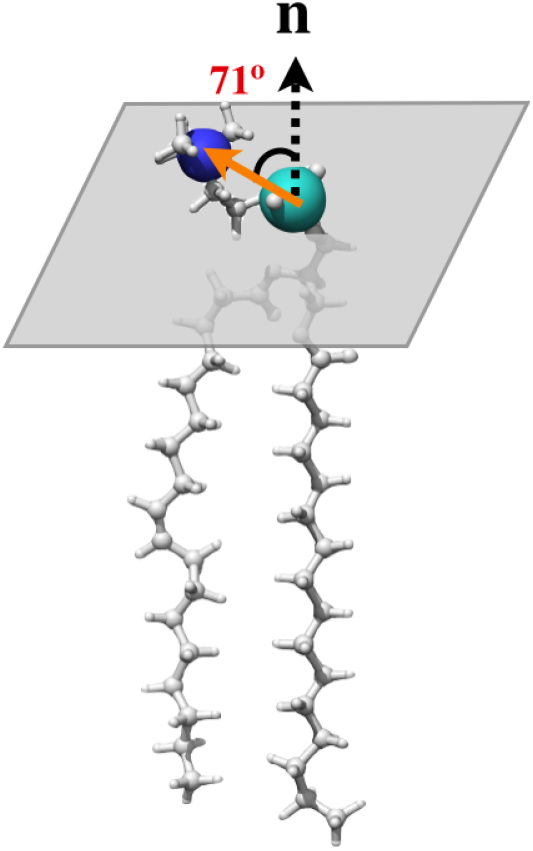
Lipid headgroup dipole in a POPC lipid (orange vector). Phosphorus and nitrogen atoms are represented by cyan and blue spheres, respectively. The headgroup dipole makes 71° with respect to the bilayer normal, making them essentially parallel to bilayer surface.

**Fig. 2.**
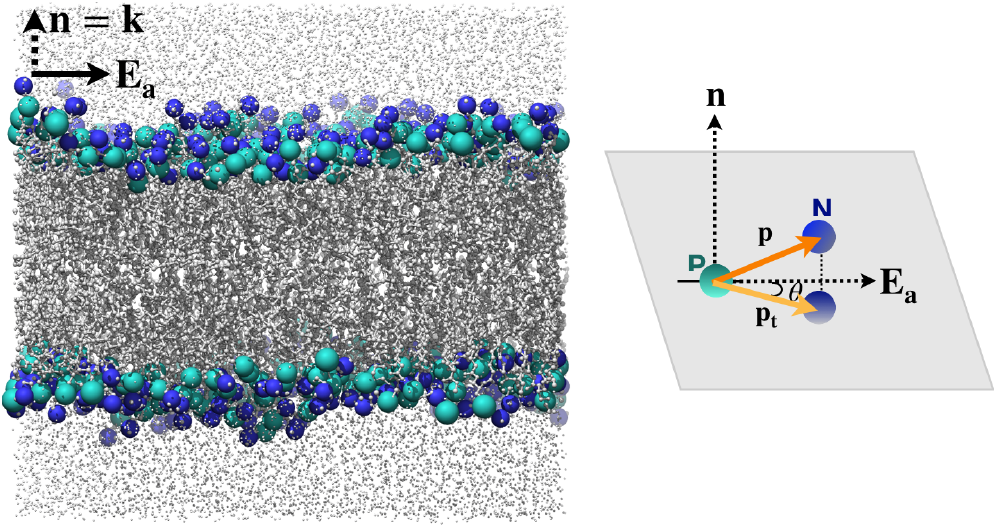
Simulated atomistic system showing the POPC lipid bilayer subjected to lateral electric field (**E**_*a*_). Phosphorus and nitrogen atoms of lipid headgroups are represented by cyan and blue colored spheres, respectively. Lipid acyl chains are shown in gray color and water molecules are represented by light gray beads on both sides of bilayer.

## Method

We adopted two approaches to quantify the in-plane polarization and the in-plane flexoelectric coefficient in lipid bilayers. We simulated all-atom flat bilayers with CHARMM36 and coarse-grained lipid vesicles with CG-MARTINI force field [27, 28]. The two systems were subjected to electric fields and the dipole reorientation and effective in-plane polarization were computed. These predictions were then used with the theoretical formulation proposed in [25] to estimate the in-plane flexoelectric coefficient. We used four different lipid types, 1-palmitoyl-2-oleoyl-sn-glycerol-3-phosphocholine (POPC), 1,2-dioleoyl-sn-glycero-3 phosphocholine (DOPC), 1,2-dipalmitoyl-sn-glycero-3-phosphocholine (DPPC) and 1-palmitoyl-2-oleoyl-sn-glycero-3-phosphoethanolamine (POPE), to capture the effect of lipid properties on the electromechanical response and the in-plane flexoelectric coefficient.

Flat bilayers were created using CHARMM-GUI [29, 30]. Each leaflet was comprised of 128 lipids and the system was solvated by 50 water molecules per lipid. The bilayers were subjected to electric fields oriented parallel to the bilayer surface and were simulated for at least 300 ns. Lipid vesicles of radius 15 nm were created in CHARMM-GUI Martini Maker. The vesicles were made of ~4500 and ~3000 lipids in the outer and the inner leaflets. More than 500,000 water molecules were used to solvate the vesicles. Each lipid vesicle was simulated for 1 *μ*s and the simulation data from the first 500 ns was discarded from the sampling analysis. The orientation of lipid head group dipoles in both the flat bilayers and the lipid vesicles were computed using an in-house code developed in MATLAB. Further details of the systems and the simulation protocols are provided in the SI.

## Results

### Predictions from flat bilayer systems

Figure 3(a) shows the effect of electric field on the dipole reorientation in the POPC bilayer. In fig. 3 (a), the x-axis is the angle (*θ*) that a dipole vector projected on to the tangent plane makes with the electric field. Thus, the angle is 0° when the vector is in the direction of the electric field and is 180° when the vector is oriented opposite to the electric field. The y-axis shows the probability density of the lipid dipoles to be oriented in the *θ* direction in the entire bilayer.

**Fig. 3.**
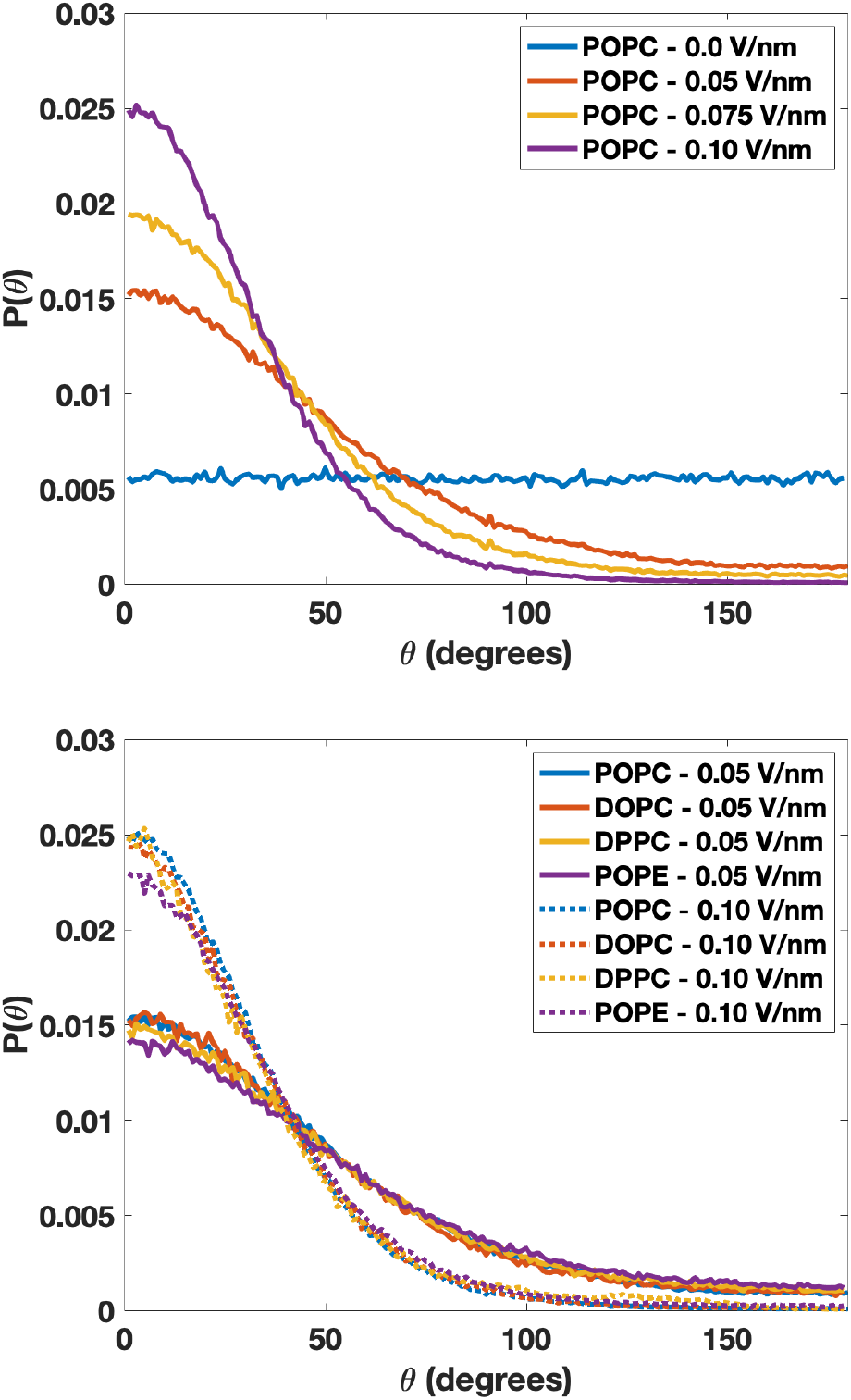
Electric field induced dipole reorientation in flat bilayers (**a**) Lateral angle probability distribution function (*P* (*θ*)) for POPC lipids when subjected to different lateral electric fields. When no electric field is applied, distribution is uniform. When lateral electric field is applied, lipid headgroup dipole vectors begin to orient along the direction of the applied electric field. The reorientation is proportional to the magnitude of the applied electric field. (**b**) *P* (*θ*) for POPC, DOPC, DPPC and POPE lipids at 0.05 V/nm and 0.10 V/nm electric field values. All four lipids show similar reorientaion of the headgroup dipoles.

In the absence of electric field, the dipoles are free to rotate in the tangent plane and hence all the orientations have equal probability (blue curve). However, when subjected to increasing electric fields, lipid dipoles begin to preferentially orient along the direction of the electric field. This increases the probability of finding a dipole vector at smaller angles with respect to the electric field direction, resulting in a rise in the *P* (*θ*) curves at smaller angles and a decline at higher angles. Once the electric field reaches 0.1 V/nm, a majority of the dipoles are aligned with the electric field.

To quantify the effect of lateral electric field on different lipid types, we simulate three additional flat bilayers made of DOPC, DPPC and POPE lipids. While POPC has one unsaturated acyl chain, DOPC and DPPC have two and zero unsaturated acyl chains, respectively. POPE lipids, on the other hand, have a different head group compared to the PC lipids. The POPE head group dipoles make an angle of 77° with respect to the bilayer normal (compared to 71° for POPC lipids). Figure 3(b) shows the probability function for the three lipids at 0.05 V/nm and 0.10 V/nm electric fields. Despite their structural differences, the dipole vectors of all three lipids show similar orientation changes as POPC for the applied electric fields.

We next used the dipole orientations to compute the in-plane polarization density (per unit area) and the in-plane flexoelectric coefficient for the four lipid systems. To compute the local in-plane polarization density in a leaflet, we divide the projected dipole vector by the area of the lipid headgroup. We average this over the two leaflets of the bilayer and sum up the two contributions to compute the resulting in-plane polarization density for the bilayer. Thus, the in-plane polarization density **π** is given by

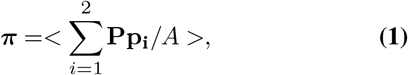

where *i* = {1, 2} accounts for the two leaflets, **P** = **I** − **n_i_** ⊗ n_i_ = **I** − **k** ⊗ **k** is the projection tensor, **p_i_** is lipid dipole vector in a leaflet, *A* is the area per lipid, and <> represents the averaging performed over the bilayer. Figure 4 shows the magnitude of **π** for the four lipid systems. The polarization density increases with an increase in the magnitude of the applied electric field as more lipids get oriented in the direction of the applied electric field. The four lipid simulations yield similar estimates.

**Fig. 4.**
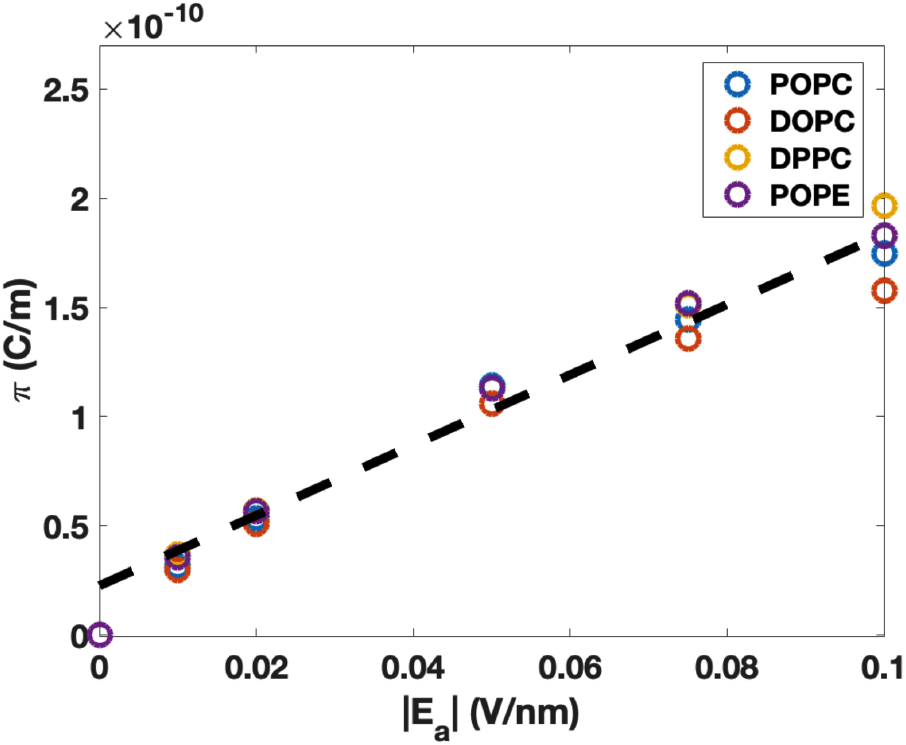
Magnitude of the in-plane polarization density (π = |**π**|) as a function of the applied electric field predicted from flat bilayer simulations. Higher electric field leads to higher dipole reorientation and hence, higher π values. A linear fit to the data (black dashed curve) yields an estiamte of the in-plane flexoelectric coefficient. Collectively, the four lipids yield an average value of *D* ≈ 5.77 × 10^17^ Nm/C^2^.

Next, we use the estimated in-plane polarization density to compute the in-plane flexoelectric coefficient for the new electromechanical theory with an in-plane polarization field. For such a system, membrane energy is given by [25]

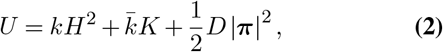

where the first two terms are the classic Helfrich-Canham energy that models the mechanical response of bilayers [31–34] and the third term is the flexoelectric term that depends on the in-plane flexoelectric coefficient *D* and the in-plane polarization density **π**. As shown in [25], equilibrium conditions yield the equation

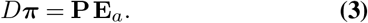

We rearrange the above equation to obtain the relation

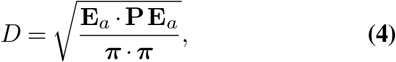

where we have used the identity **P**^*T*^**P** = **P**. For the flat bilayer systems, **PE**_*a*_ = **E**_*a*_. Therefore, the expression for the in-plane flexoelectric coefficient reduces to

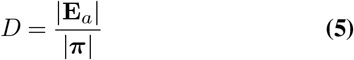

Thus, the in-plane flexoelectric coefficient is the reciprocal of the slope of the π(|**E**_*a*_|) curves. The linear fit (black dashed curve) in Figure 4 yields an estimate of *D* ≈ 5.77 × 10^17^ Nm/C^2^ for the four simulated lipid types.

### Predictions from vesicle systems

To verify the estimate obtained from the flat bilayers systems, we constructed coarse-grained lipid vesicles in Martini. In particular, we wanted to determine if the curvature of the bilayers would impact the predictions of the in-plane flexoelectric coefficient. We constructed spherical vesicles for the four lipid types and subjected them to an electric field of 0.025 V/nm. Figure 5 shows the deformed state of a POPC vesicle in the presence of the applied electric field. The inset shows a schematic of the lipid dipole vector, the applied electric field vector and their projected components on to the tangential plane of the bilayer surface.

**Fig. 5.**
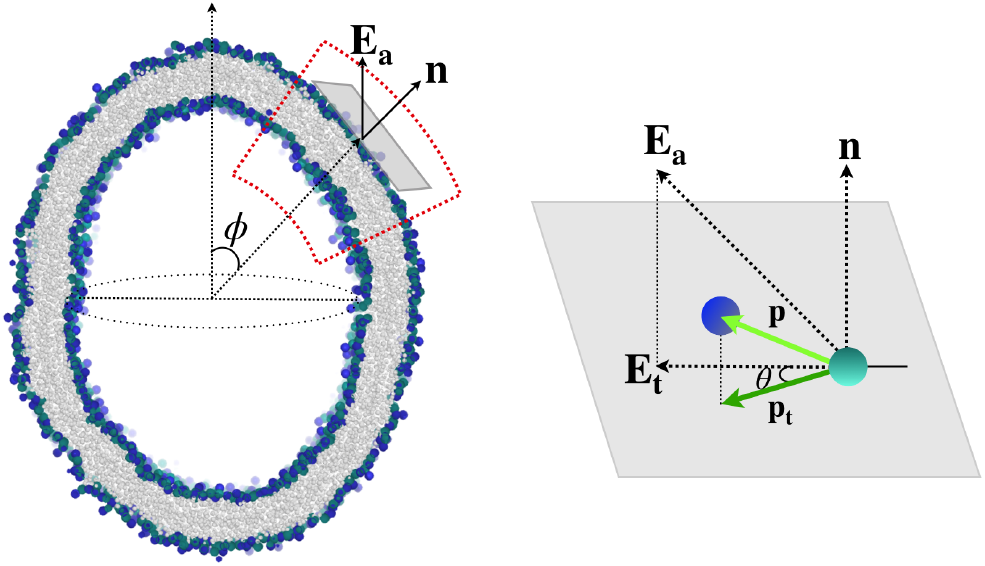
Coarse-grained POPC vesicle in the presence of an applied electric field (0.025 V/nm). The initial spherical vesicle turns into a prolate ellipsoid in the presence of the electric field. The electric field-induced reorientation of the dipole vector is captured via the angle *θ* between the projected dipole vector **p_t_** and the projected electric field vector **E_t_** onto the tangent plane of the vesicle surface.

Figure 6(a) shows the probability density *P* (*θ*) of the projected dipoles to align with the projected electric field **E**_*t*_. As before, in the absence of the electric field all angles have equal probability yielding the flat curves for the four lipid systems. In the presence of the electric field, the dipoles become oriented in the direction of **E**_*t*_ resulting in an increase in *P* (*θ*) at smaller angles and a decline at larger angles. The trend is consistent for the four lipid types. Next, we use eqn. 1 to compute the in-plane polarization density from the projected dipole vectors in the bilayer. Figure 6(b) shows |**π**| as we move along a meridian from the north pole to the south pole of the simulated vesicle. We present the data for 20° ≤ *ϕ* ≤ 160° due to high numerical error encountered in the close vicinity of the two poles because of the vanishing denominator in eqn. 1. Figure 6(b) shows that around the two poles, the magnitude of in-plane polarization density is small. This is so because **E**_*t*_ is small and hence, the dipoles continue to have their rotational freedom. Around the equatorial plane (*ϕ* ≈ 90°), the in-plane polarization density achieves its maximum value. In this region, **E**_*t*_ is maximum, which leads to the maximum alignment of lipid dipoles.

**Fig. 6.**
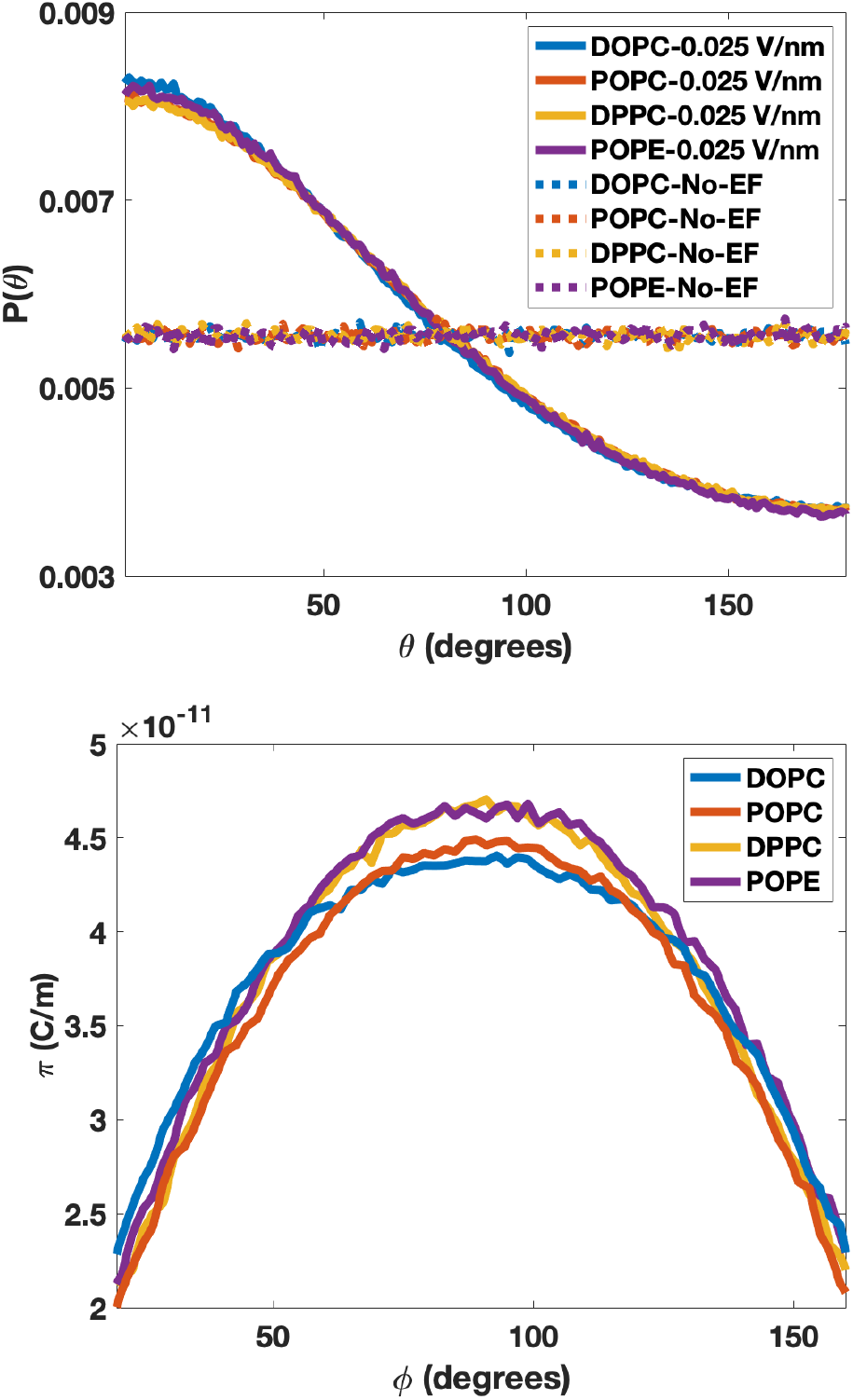
Predictions from the vesicle simulations. (**a**) *P* (*θ*) plots for POPC, DOPC, DPPC and POPE lipids in the absence and presence of 0.025V/nm electric field. In the absence of electric field, *P* (*θ*) plots are flat as dipoles undergo free rotation. In the presence of electric field, dipoles undergo reorientation and *P* (*θ*) increases for smaller angles and decreases for larger angles. The plots show similar trends for all the four lipid types. (**b**) The magnitude of in-plane polarization density as we move along a meridian from the north pole to the south pole of the deformed vesicle (20° ≤ *ϕ* ≤ 160°). The magnitude increases as we away from the poles towards the equator, in proportion to the magnitude of the projected electric field onto the tangent plane and the dipole reorientation.

Finally, we use eqn. 4 to calculate the in-plane flexoelectric coefficient for the four lipid systems. Figure 7 shows the predicted values as a function of the polar angle. The plots yield an average value of 5.5 × 10^17^ Nm/C^2^ for the in-plane flexoelectric coefficient for the different lipid types. These estimates are in very good agreement with the earlier estimate obtained from the atomistic simulations. Since both the flat bilayer and the vesicle geometries yield similar estimates, *D* is independent of membrane geometry, and hence, is purely a material property.

**Fig. 7.**
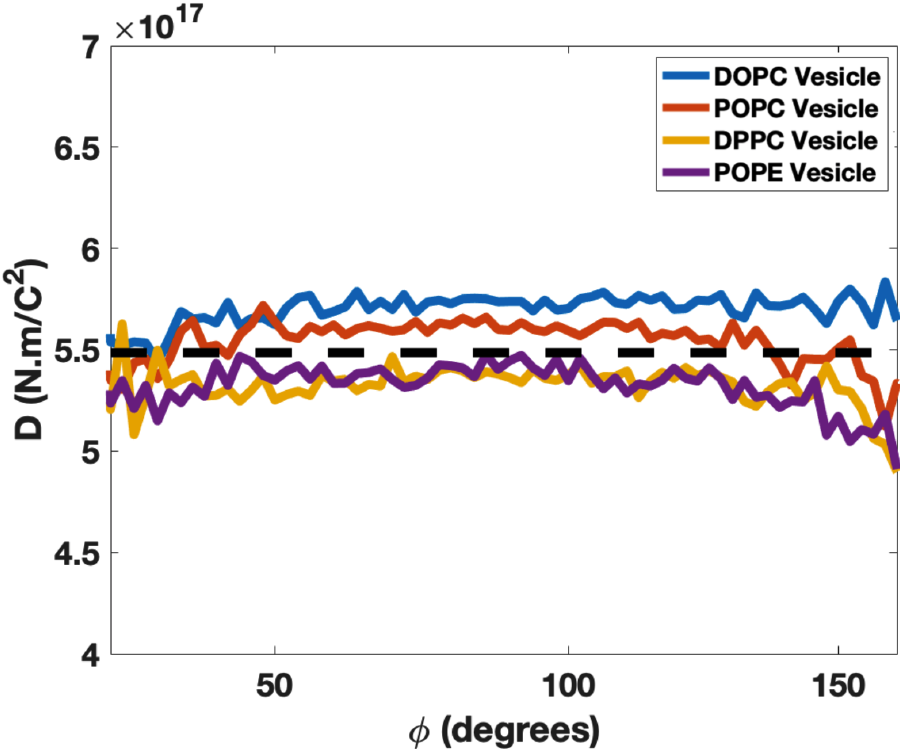
The predicted in-plane flexoelectric coefficient obtained from the vesicle simulations. These simulations yield an average value of 5.5 × 10^17^ Nm/C^2^ for the four lipid types (black dashed curve).

## Conclusions

In this study, we performed atomistic and coarse-grained molecular dynamics simulations to quantify the propensity of lipid bilayers to undergo in-plane polarization in the presence of applied electric field. We used the numerical predictions to estimate the in-plane flexoelectric coefficient required to model the electromechanical response of lipid bilayers. Our results show that the in-plane flexoelectric coefficient *D* ≈ 5.5 × 10^17^ Nm/C^2^ for the four simulated lipid types. Despite the variations in the acyl chain saturation level and the head groups structure, the four lipids show a comparable in-plane polarization.

Our study provides the first molecular proof of the in-plane polarization undergone by lipid bilayers in the presence of an applied electric field. The study shows that the bilayers undergo significant in-plane polarization, suggesting that in-plane polarization should be accounted for in analyzing the electromechanical response of bilayers in the presence of electric fields with tangential component. Our simulations also show that the angle dipoles make with the bilayer normal undergo minimal changes in the presence of applied electric field (Fig. S1) suggesting that in-plane polarization should be the key factor determining the electromechanical response of bilayers. Furthermore, our study shows that the *D* estimate is independent of the bilayer geometry, confirming the assumption of a constant material property invoked in the new electromechanical theory [25]. Estimation of *D* overcomes a major roadblock that would allow future studies to numerically quantify the electromechanical response of membranes and validate modeling predictions with experimental data.

## ACKNOWLEDGEMENTS

A.A. acknowledges support from NSF Grants CMMI-1727271 and CMMI-1931084. The authors acknowledge the use of the Opuntia, Sabine, and Carya clusters to perform the simulations and the advanced technical support from the Research Computing Data Core at UH to carry out the research presented here.

## Notes

### Competing Interest Statement

The authors have declared no competing interest.

## References

1. Richard Nuccitelli. A role for endogenous electric fields in wound healing. Current topics in developmental biology, 58(2):1–26, 2003.

2. Guangping Tai, Michael Tai, and Min Zhao. Electrically stimulated cell migration and its contribution to wound healing. Burns & trauma, 6(1), 2018.

3. Entong Wang, Min Zhao, John V Forrester, and Colin D McCaig. Electric fields and map kinase signaling can regulate early wound healing in lens epithelium. Investigative ophthalmology & visual science, 44(1):244–249, 2003.

4. Jason T Sengel and Mark I Wallace. Imaging the dynamics of individual electropores. Proceedings of the National Academy of Sciences, 113(19):5281–5286, 2016.

5. Mayya Tokman, Jane HyoJin Lee, Zachary A Levine, Ming-Chak Ho, Michael E Colvin, and P Thomas Vernier. Electric field-driven water dipoles: nanoscale architecture of electroporation. PloS one, 8(4):e61111, 2013.

6. Andraž Polak, Mounir Tarek, Matija Tomšič, Janez Valant, Nataša Poklar Ulrih, Andrej Jamnik, Peter Kramar, and Damijan Miklavčič. Electroporation of archaeal lipid membranes using md simulations. Bioelectrochemistry, 100:18–26, 2014.

7. Rainer A Böckmann, Bert L De Groot, Sergej Kakorin, Eberhard Neumann, and Helmut Grubmüller. Kinetics, statistics, and energetics of lipid membrane electroporation studied by molecular dynamics simulations. Biophysical journal, 95(4):1837–1850, 2008.

8. Lucie Delemotte and Mounir Tarek. Molecular dynamics simulations of lipid membrane electroporation. The Journal of membrane biology, 245(9):531–543, 2012.

9. Gunter A Hofmann, SB Dev, S Dimmer, and GS Nanda. Electroporation therapy: a new approach for the treatment of head and neck cancer. IEEE Transactions on Biomedical Engineering, 46(6):752–759, 1999.

10. Natanel Jourabchi, Kourosh Beroukhim, Bashir A Tafti, Stephen T Kee, and Edward W Lee. Irreversible electroporation (nanoknife) in cancer treatment. Gastrointestinal Intervention, 3 (1):8–18, 2014.

11. Anita Gothelf, Lluis M Mir, and Julie Gehl. Electrochemotherapy: results of cancer treatment using enhanced delivery of bleomycin by electroporation. Cancer treatment reviews, 29(5): 371–387, 2003.

12. William E Brownell, Charles R Bader, Daniel Bertrand, and Yves De Ribaupierre. Evoked mechanical responses of isolated cochlear outer hair cells. Science, 227(4683):194–196, 1985.

13. Robert M Raphael, Aleksander S Popel, and William E Brownell. A membrane bending model of outer hair cell electromotility. Biophysical journal, 78(6):2844–2862, 2000.

14. Ben Harland, Wen-han Lee, William E Brownell, Sean X Sun, and Alexander A Spector. The potential and electric field in the cochlear outer hair cell membrane. Medical & biological engineering & computing, 53(5):405–413, 2015.

15. Rumiana Dimova, Natalya Bezlyepkina, Marie Domange Jordö, Roland L Knorr, Karin A Riske, Margarita Staykova, Petia M Vlahovska, Tetsuya Yamamoto, Peng Yang, and Reinhard Lipowsky. Vesicles in electric fields: some novel aspects of membrane behavior. Soft Matter, 5(17):3201–3212, 2009.

16. Rumiana Dimova. Recent developments in the field of bending rigidity measurements on membranes. Advances in colloid and interface science, 208:225–234, 2014.

17. Petia M Vlahovska. Nonequilibrium dynamics of lipid membranes: Deformation and stability in electric fields. In Advances in Planar Lipid Bilayers and Liposomes, volume 12, pages 101–146. Elsevier, 2010.

18. Jia Zhang, Jeffrey D Zahn, Wenchang Tan, and Hao Lin. A transient solution for vesicle electrodeformation and relaxation. Physics of Fluids, 25(7):071903, 2013.

19. AT Todorov, AG Petrov, and JH Fendler. First observation of the converse flexoelectric effect in bilayer lipid membranes. The Journal of Physical Chemistry, 98(12):3076–3079, 1994.

20. Alexander G Petrov. Flexoelectricity of model and living membranes. Biochimica et Biophysica Acta (BBA)-Biomembranes, 1561(1):1–25, 2002.

21. M Kummrow and W Helfrich. Deformation of giant lipid vesicles by electric fields. Physical Review A, 44(12):8356, 1991.

22. Zhanchun Tu, Jixing Liu, Yuzhang Xie, and Zhong-can Ou-yang. Geometric Methods in Elastic Theory of Membranes in Liquid Crystal Phases, volume 2. World Scientific, 2017.

23. P Mohammadi, LP Liu, and Pradeep Sharma. A theory of flexoelectric membranes and effective properties of heterogeneous membranes. Journal of Applied Mechanics, 81(1), 2014.

24. Ling-Tian Gao, Xi-Qiao Feng, Ya-Jun Yin, and Huajian Gao. An electromechanical liquid crystal model of vesicles. Journal of the Mechanics and Physics of Solids, 56(9):2844–2862, 2008.

25. David Steigmann and Ashutosh Agrawal. Electromechanics of polarized lipid bilayers. Mathematics and Mechanics of Complex Systems, 4(1):31–54, 2016.

26. Niloufar Abtahi, Lila Bouzar, Nadia Saidi-Amroun, and Martin Michael Müller. Flexoelectric fluid membrane vesicles in spherical confinement. EPL (Europhysics Letters), 131(1): 18001, jul 2020. doi: 10.1209/0295-5075/131/18001.

27. Jing Huang, Sarah Rauscher, Grzegorz Nawrocki, Ting Ran, Michael Feig, Bert L de Groot, Helmut Grubmüller, and Alexander D MacKerell. Charmm36m: an improved force field for folded and intrinsically disordered proteins. Nature methods, 14(1):71–73, 2017.

28. Siewert J Marrink, H Jelger Risselada, Serge Yefimov, D Peter Tieleman, and Alex H De Vries. The martini force field: coarse grained model for biomolecular simulations. The journal of physical chemistry B, 111(27):7812–7824, 2007.

29. Jumin Lee, Xi Cheng, Jason M Swails, Min Sun Yeom, Peter K Eastman, Justin A Lemkul, Shuai Wei, Joshua Buckner, Jong Cheol Jeong, Yifei Qi, et al. Charmm-gui input generator for namd, gromacs, amber, openmm, and charmm/openmm simulations using the charmm36 additive force field. Journal of chemical theory and computation, 12(1):405–413, 2016.

30. Yifei Qi, Helgi I Ingólfsson, Xi Cheng, Jumin Lee, Siewert J Marrink, and Wonpil Im. Charmm-gui martini maker for coarse-grained simulations with the martini force field. Journal of chemical theory and computation, 11(9):4486–4494, 2015.

31. Peter B Canham. The minimum energy of bending as a possible explanation of the biconcave shape of the human red blood cell. Journal of theoretical biology, 26(1):61–81, 1970.

32. Wolfgang Helfrich. Elastic properties of lipid bilayers: theory and possible experiments. Zeitschrift für Naturforschung C, 28(11-12):693–703, 1973.

33. James T Jenkins. The equations of mechanical equilibrium of a model membrane. SIAM Journal on Applied Mathematics, 32(4):755–764, 1977.

34. David J Steigmann. A model for lipid membranes with tilt and distension based on three-dimensional liquid crystal theory. International Journal of Non-Linear Mechanics, 56:61–70, 2013.

