## Supplemental Document for "Quantification of in-plane flexoelectricity in lipid bilayers"

**1 Department of Mechanical Engineering, University of Houston, Houston, TX 77204**

**Corresponding authors:**
*****

### Molecular dynamics simulation protocol

#### Lipid bilayer:

All-atom models of DOPC,POPC,DPPC and POPE bilayer systems were created using CHARMM-GUI [1]. 128 lipids per leaflets were used to create the bilayer. 50 water molecules per lipids were added to the bilayer on both side of the bilayer. We used CHARMM36m [2] force field to simulate the systems in GROMACS 2018 [3]. Steepest descent algorithm was used for energy minimization. Energy minimization was performed until maximum force in the system was below 700 KJ/mol.nm. We performed three stages NVT equilibration and three stages of NPT equilibration for a total of 500 picoseconds prior to the NPT production runs. Berendsen thermocouple [4] with 1.0 ps time constant and Semiisotropic Berendsen pressure couple with 5.0 ps were used throughout the equilibration. Production runs were performed with Nosé-Hoover thermocouple [5] with 1.0 ps time constant and semiisotropic Parrinello-Rahman pressure couple [6] with 5.0 ps time constant. Pressure of the systems were maintained at $10^5$ bar with 4.5 $\times 10^5$ bar $^{-1}$ compressibility throughout the simulation. All simulations were performed at temperatures above phase transition temperature of lipids. Simulation temperature of all systems are provided in Table 1. Verlet cut-off scheme was used throughout the simulation. Van der Waals interactions were cut-off at 1.2 nm. We used force-switch vdw-modifier at 1.0 nm. Real space coulombic interactions were cut-off at 1.2 nm and reciprocal space interactions were calculated using PME [7]. Hydrogen bonds were constrained using LINCS algorithm [8]. Systems with external electric field intensities were modeled by applying electric fields in Y-direction, parallel to the bilayer surface. Production runs were performed for at least 300 ns. first 150 ns of simulations were ignored for equilibration. Last 150 ns with 20 ps interval were used for sampling. Simulation systems with 0.10 V/nm used bilayer systems with 48 lipids per leaflet instead of 128 lipids since 128 lipid systems were buckling with 0.10 V/nm external lateral electric field application. These systems were simulated for 600 ns after the initial equilibration.

#### Lipid vesicle:

Lipid vesicles of radius 15 nm were created in CHARMM-GUI Martini Maker [9]. POPC, DOPC, DPPC and POPE lipids were chosen to create the vesicle. 15 nm radius vesicle has $\sim$ 7500 lipids with 4500 and 3000 lipids in outer and inner leaflet. More than 500,000 water molecules were used to solvate larger vesicles. Since these lipids were neutral, no additional salt was added into the solution. Steepest descent energy minimization performed for 10000 steps. Verlet cut-off scheme was used with 0.005 buffer tolerance. Reaction field method was used to compute the coulombic interactions with dielectric constant beyond the 1.1 nm cut-off set at 2.5. Van derWaal's interactions were cut-off at 1.1 nm and potential shift verlet modifier was used as vdw-modifier. Systems were initially equilibrated for 100 ns with 0.001 fs time increment. PO4 bead of each lipids were constrained to ensure that vesicles maintain the spherical geometry during the initial equilibration. V-rescale thermocouple [10] was used to maintain the system temperature at 323.15 K, with time constant of 1.0 ps. In addition, berendsen isotropic pressure coupling [4] with 5.0 ps time constant and $4.5 \times 10^{-5}$ bar $^{-1}$ compressibility was used to maintain the pressure at $10^5$ bar. 2 fs timestep was used throughout this equilibration. This equilibration was followed by another four equilibration steps, in which constraints were gradually removed. The production run were performed for 1 microsecond with 0.010 ps timestep. First 500 ns simulation data was discarded for sampling. When external 0.025 V/nm electric field was applied in Z-direction throughout the system when polarization simulations were performed. Further details of systems and their simulation protocol are provided in Table 2.

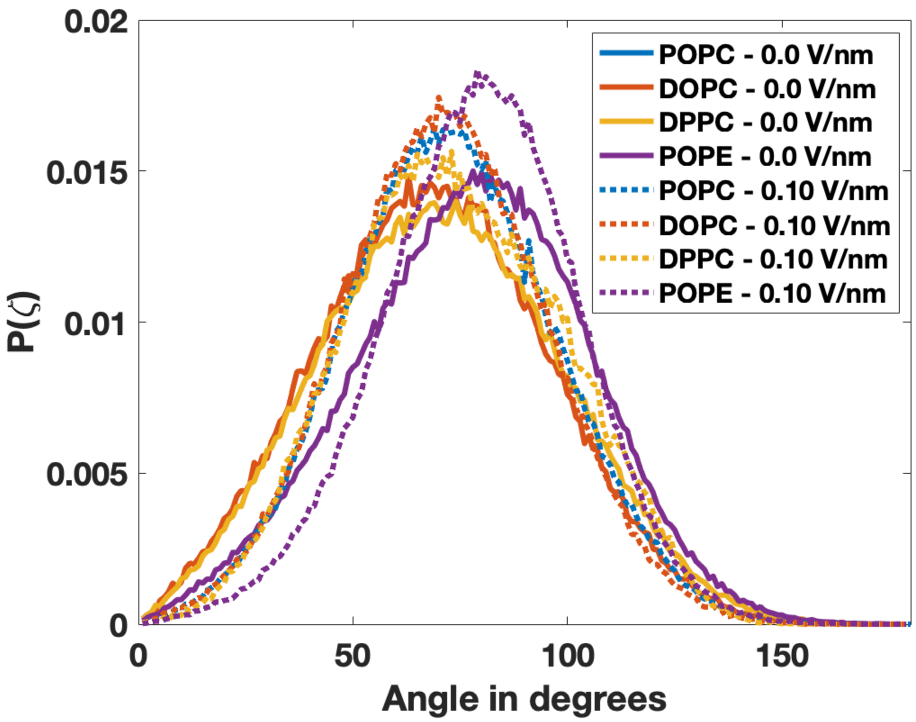


Figure S1: Normal angle probability density distribution of POPC,DOPC,DPPC and POPE lipids upon different 0.0 V/nm and 0.10 V/nm external lateral electric field. $\zeta$ is the angle that dipole vector makes with the bilayer normal ($\mathbf{k}$). Dipole prefers to orient close to 90 degree with bilayer normal when electric field applied. PN dipole of POPE lipids lie flatter to the bilayer surface than PC lipids.

#### Lipid vesicle calculation:

Coordinates of PO4 beads of lipids were taken for computation of surface normal. PO4 and NC3 beads of PC lipids were chosen to define lipid headgroup dipole vector. NH3 beads were chosen instead of NC3 for PE lipids. Area per lipid of outer ($A_{out}$) and inner ($A_{in}$) leaflets of a vesicle is computed as follows.

$$A_j^i = \frac{S_j^i}{N_j^{Lipids}} \qquad [S1]$$

where $\{i\}$ represents the inner or outer leaflet in $j^{th}$ frame. $S_j^i$ is the total surface area of the inner or outer leaflet in $j^{th}$ frame. $N_{Lipids}^i$ is the total number of lipids in inner or outer leaflet.

Surface normals at each PO4 beads in a single frame were calculated using 'normal.m' available in MATLAB. $\mathbf{n_{jk}^i}$ is the surface normal vector at $k^{th}$ PO4 bead in $j^{th}$ frame in the inner or outer leaflet. Similarly, $\mathbf{d_{jk}^i}$ represents the lipid dipole vector of that particular lipid. Once, we compute the surface normal and dipole vector, we can compute in-plane polarization density ($\boldsymbol{\pi}$) and D as follows.

$$\mathbf{P_{jk}^i} = I - \mathbf{n_{jk}^i} \otimes \mathbf{n_{jk}^i} \qquad [S2]$$

where $\mathbf{P_{jk}^i}$ is the projection tensor at that point.
in-plane polarization density is computed as below.

$$\boldsymbol{\pi}_{\mathbf{jk}}^{\mathbf{i}} = \frac{\mathbf{P_{jk}^i d_{jk}^i}}{A_j^i} \qquad [S3]$$

In addition, we can also compute the numerator of D.

$$D_{num}{}^{i}_{jk} = \sqrt{\mathbf{E_a}.\mathbf{P^i_{jk}}\mathbf{E_a}} \quad [S4]$$

Once, we obtain these parameters, we perform the grid-based averaging. We compute the average of in-plane polarization density and numerator of D for each $(\theta, \phi)$ grid point. After, we obtain the total in-plane polarization density as follows.

$$\boldsymbol{\pi}^{Total}(\theta, \phi) = \boldsymbol{\pi}^{in}(\theta, \phi) + \boldsymbol{\pi}^{out}(\theta, \phi) \quad [S5]$$

We can compute the numerator of D parameter for $(\theta, \phi)$ by taking average of outer and inner leaflets.

$$D^{avg}_{num}(\theta, \phi) = \frac{1}{2}(D^{in}_{num}(\theta, \phi) + D^{out}_{num}(\theta, \phi)) \quad [S6]$$

$$D(\theta, \phi) = \frac{D^{avg}_{num}(\theta, \phi)}{\sqrt{\boldsymbol{\pi}^{Total}(\theta, \phi).\boldsymbol{\pi}^{Total}(\theta, \phi)}} \quad [S7]$$

### Calculation of probability density

$P(\theta)$ is the probability density of $\theta$ angle distribution. $\theta$ angle is computed as follows.

$$\theta = acos(\frac{\mathbf{PN_t}.\mathbf{E_t}}{|\mathbf{PN_t}||\mathbf{E_t}|}) \quad [S8]$$

where $\mathbf{PN_t}$ is the projection of $\mathbf{PN}$ onto tangent plane and for a flat bilayer, tangent plane is the XY-plane. For a flat bilayer, electric field was applied along Y-direction. Therefore, $\mathbf{E_t}$ is $\mathbf{E_a}$ itself.

$$\theta = acos(\frac{\mathbf{PN_t}.\mathbf{E_a}}{|\mathbf{PN_t}||\mathbf{E_a}|}) \quad [S9]$$

After computing the $\theta$ angle distribution, histogram representation of $\theta$ angle distribution is generated with 1° angle increment. Probability density is computed by the following formula.

$$P(\theta_i) = \frac{h(\theta_i)}{N_l \times N_{sampling}} \quad [S10]$$

$$h(\theta_i) = \sum_{n_f=0}^{N_{sampling}} \sum_{n=1}^{N_l} H(\theta - (\theta_i - \delta\theta)) - H(\theta - (\theta_i + \delta\theta)) \quad [S11]$$

where $P(\theta_i)$ is the probability density corresponding to $\theta_i$ angle, $N_{sampling}$ is the total number of sampling frames in a simulation, $N_l$ is the total number of lipids in a bilayer, $h(\theta_i)$ is the total number of lipids for which $\theta$ angle is $(\theta_i - \delta\theta \leq \theta \leq \theta_i + \delta\theta).H(\theta)$ is the Heaviside function and $\delta\theta$ is the angle increment to create the bins.

| System | Lipids per leaflet | Time (ns) | Temperature (K) | EF (V/nm) |
|---|---|---|---|---|
| DPPC | 128 | 300 | 323 | 0.0 |
| DPPC | 128 | 300 | 323 | 0.01 |
| DPPC | 128 | 300 | 323 | 0.02 |
| DPPC | 128 | 300 | 323 | 0.05 |
| DPPC | 128 | 300 | 323 | 0.075 |
| DPPC | 48 | 600 | 323 | 0.10 |
| POPC | 128 | 300 | 303 | 0.0 |
| POPC | 128 | 300 | 303 | 0.01 |
| POPC | 128 | 300 | 303 | 0.02 |
| POPC | 128 | 300 | 303 | 0.05 |
| POPC | 128 | 300 | 303 | 0.075 |
| POPC | 48 | 600 | 303 | 0.10 |
| DOPC | 128 | 300 | 303 | 0.0 |
| DOPC | 128 | 300 | 303 | 0.01 |
| DOPC | 128 | 300 | 303 | 0.0.02 |
| DOPC | 128 | 300 | 303 | 0.05 |
| DOPC | 128 | 300 | 303 | 0.075 |
| DOPC | 48 | 600 | 303 | 0.10 |
| POPE | 128 | 300 | 323 | 0.0 |
| POPE | 128 | 300 | 323 | 0.01 |
| POPE | 128 | 300 | 323 | 0.0.02 |
| POPE | 128 | 300 | 323 | 0.05 |
| POPE | 128 | 300 | 323 | 0.075 |
| POPE | 128 | 600 | 323 | 0.10 |

Table 1: Summary of all-atom MD simulations of lipid bilayers. All simulations were performed above phase transition temperatures.

| System | Lipids in outer leaflet | Lipids in inner leaflet | Time (ns) | Temperature (K) | EF (V/nm) |
|---|---|---|---|---|---|
| DPPC | 4308 | 3348 | 1000 | 323 | 0.0 |
| DPPC | 4308 | 3348 | 1000 | 323 | 0.025 |
| POPC | 4263 | 3269 | 1000 | 323 | 0.0 |
| POPC | 4263 | 3269 | 1000 | 323 | 0.025 |
| POPE | 4342 | 3411 | 1000 | 323 | 0.0 |
| POPE | 4342 | 3411 | 1000 | 323 | 0.025 |
| DOPC | 4251 | 3249 | 1000 | 323 | 0.0 |
| DOPC | 4251 | 3249 | 1000 | 323 | 0.025 |

Table 2: Summary of coarse grained MARTINI MD simulations of lipid vesicles. Each type of lipid vesicles were simulated with and without electric field.

### References


1. Jumin Lee, Xi Cheng, Jason M Swails, Min Sun Yeom, Peter K Eastman, Justin A Lemkul, Shuai Wei, Joshua Buckner, Jong Cheol Jeong, Yifei Qi, et al. Charmm-gui input generator for namd, gromacs, amber, openmm, and charmm/openmm simulations using the charmm36 additive force field. *Journal of chemical theory and computation*, 12(1):405–413, 2016.

2. Jing Huang, Sarah Rauscher, Grzegorz Nawrocki, Ting Ran, Michael Feig, Bert L de Groot, Helmut Grubmüller, and Alexander D MacKerell. Charmm36m: an improved force field for folded and intrinsically disordered proteins. *Nature methods*, 14(1):71–73, 2017.

3. Mark James Abraham, Teemu Murtola, Roland Schulz, Szilárd Páll, Jeremy C Smith, Berk Hess, and Erik

Lindahl. Gromacs: High performance molecular simulations through multi-level parallelism from laptops to supercomputers. *SoftwareX*, 1:19–25, 2015.

4. Herman JC Berendsen, JPM van Postma, Wilfred F van Gunsteren, ARHJ DiNola, and Jan R Haak. Molecular dynamics with coupling to an external bath. *The Journal of chemical physics*, 81(8):3684–3690, 1984.

5. Shuichi Nosé. A unified formulation of the constant temperature molecular dynamics methods. *The Journal of chemical physics*, 81(1):511–519, 1984.

6. Michele Parrinello and Aneesur Rahman. Polymorphic transitions in single crystals: A new molecular dynamics method. *Journal of Applied physics*, 52(12):7182–7190, 1981.

7. Tom Darden, Darrin York, and Lee Pedersen. Particle mesh ewald: An n log (n) method for ewald sums in large systems. *The Journal of chemical physics*, 98(12):10089–10092, 1993.

8. Berk Hess, Henk Bekker, Herman JC Berendsen, and Johannes GEM Fraaije. Lincs: a linear constraint solver for molecular simulations. *Journal of computational chemistry*, 18(12):1463–1472, 1997.

9. Yifei Qi, Helgi I Ingólfsson, Xi Cheng, Jumin Lee, Siewert J Marrink, and Wonpil Im. Charmm-gui martini maker for coarse-grained simulations with the martini force field. *Journal of chemical theory and computation*, 11(9):4486–4494, 2015.

10. Giovanni Bussi, Davide Donadio, and Michele Parrinello. Canonical sampling through velocity rescaling. *The Journal of chemical physics*, 126(1):014101, 2007.
